# Transcriptomic profiling of retinal cells reveals a subpopulation of microglia/macrophages expressing Rbpms and Spp1 markers of retinal ganglion cells (RGCs) that confound identification of RGCs

**DOI:** 10.1101/2023.01.23.525216

**Authors:** William C. Theune, Ephraim F. Trakhtenberg

## Abstract

Analysis of retinal ganglion cells (RGCs) by scRNA-seq is emerging as a state-of-the-art method for studying RGC biology and subtypes, as well as for studying the mechanisms of neuroprotection and axon regeneration in the central nervous system (CNS). Rbpms has been established as a pan-RGC marker, and Spp1 has been established as an αRGC type marker. Here, we analyzed by scRNA-seq retinal microglia and macrophages, and found Rbpms+ and Spp1+ subpopulations of retinal microglia/macrophages, which pose a potential pitfall in scRNA-seq studies involving RGCs. We performed comparative analysis of cellular identity of the presumed RGC cells isolated in recent scRNA-seq studies, and found that Rbpms+ and Spp1+ microglia/macrophages confounded identification of RGCs. We also provide solutions for circumventing this potential pitfall in scRNA-seq studies, by including in RGC and αRGC selection criteria other pan-RGC and αRGC markers.

## INTRODUCTION

The Pten tumor suppressor gene is one of the most potent gene-regulators of CNS axon regeneration. Pten suppresses regeneration through inhibition of the mTOR pathway, and Pten knockout (KO) and knockdown (KD) have been shown to promote various extents of axon regeneration, after optic nerve crush (ONC) injury in mice, in the α and intrinsically photosensitive (ip) subset of retinal ganglion cells (RGCs)^1-4^. In order to understand the effect of Pten inhibition on the transcriptomes of RGCs that respond by regenerating axons of any length, recent studies by Li et al.^5^ and by Jacobi et al.^6^ isolated the RGCs with Pten KO that regenerated axons at least short-distance (1.5 mm) and profiled them by scRNA-seq. Here, we performed comparative analysis of cellular identity of the presumed RGC cells isolated in these and other relevant studies, and reveal the existence of Rbpms+ and Spp1+ subpopulations of microglia/macrophages, which confounded identification of RGCs in scRNA-seq studies.

## RESULTS

We report that, the pan-RGC marker Rbpms is also expressed in a subset of microglia and macrophages (which infiltrate the CNS after injury). Therefore, Rbpms alone is insufficient for identifying RGCs in scRNA-seq analysis. For example, in the study by Li et al.^5^, Pten KO-treated RGCs were selected from within scRNA-seq-profiled retinal cells based on the expression of at least one RGC marker (either Rbpms, Thy1, Slc17a6, or Pou4f1-3), rather than co-expression of multiple markers. This led to ∼94% of the cells selected as presumed RGCs to be identified as microglia/macrophages by retrospect analyses (see below), which is consistent with top enriched genes being microglia/macrophage markers and the top enriched gene-networks being microglia/macrophage-related (e.g., leukocyte migration, lymphocyte mediated immunity), as reported in that study^5^. Thus, in order to determine that a cell is an RGC, several of the well-established pan-RGC (e.g., Slc17a6 and Sncg) and neuronal markers (e.g., Tubb3 and Syn1) need to be co-expressed in a cell.

To demonstrate this potential pitfall of relying only on expression of a single RGC marker rather than co-expression of several markers in scRNA-seq studies, we performed comparative analysis on the cells with Pten KO labeled as RGCs in the studies by Li et al.^5^ and Jacobi et al.^6^. We also included in the analysis scRNA-seq-profiled microglia/macrophages, which we isolated from retinas and optic nerves, and separated based on the expression of Rbpms (Fig. 1A). The retinal microglia/macrophage Rbpms+ subpopulation represented 12% of all retinal microglia/macrophages. Rbpms detection was not a consequence of a piece of broken RGC sticking to microglia/macrophage (a doublet), because we found that 5% of all optic nerve microglia/macrophages also expressed Rbpms, and no RGC soma is found in the optic nerve. Furthermore, no other RGC markers were found in either subpopulation of microglia/macrophages (Fig. 1A-E). The Rbpms+ subpopulation overall represented 6.6% of all retinal and optic nerve microglia/macrophage cells.

**Figure 1.**
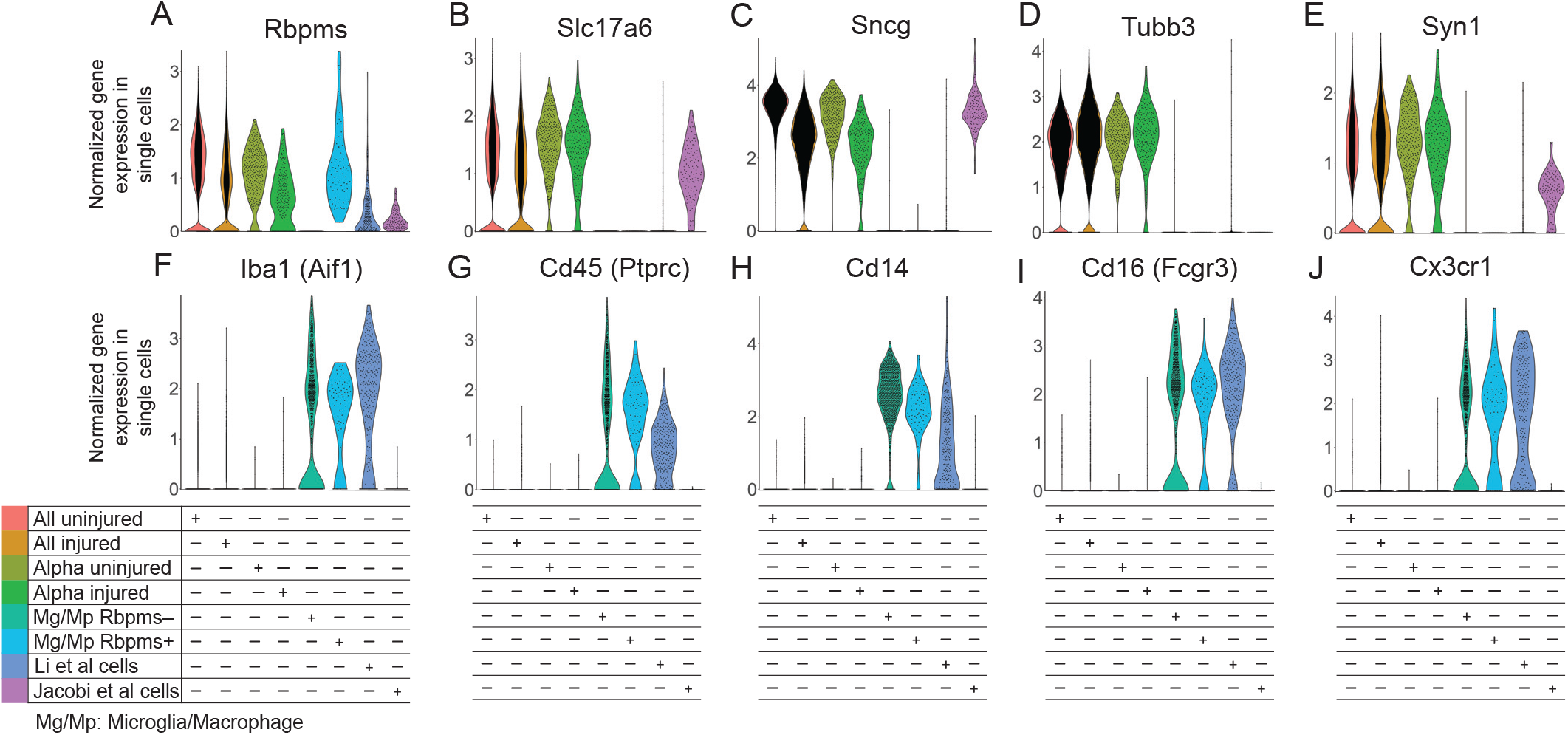
Violin plots of pan-RGC and microglia/macrophage gene markers in single cells from different conditions. (**A**-**E**) Pan-RGC (*A-C*) and neuronal-specific (*D-E*) gene markers are enriched in Tran et al. (2019) uninjured and injured RGCs, including in the αRGC subset, but with an exception for Rbpms (*A*) are not expressed in the microglia/macrophages (Mg/Mp). Pan-RGC (*A-C*) and neuronal-specific (*D-E*) gene markers are also enriched in axon-regenerating RGCs isolated by Jacobi et al. (2022), but are absent from the majority of presumed RGC cells isolated by Li et al. (2022), with an exception for Rbpms (*A*) which is also enriched in the Rbpms+ subpopulation of microglia/macrophages. Tubb3 (*D*) is enriched only in uninjured and injured untreated RGCs. (**F**-**J**) Microglia/macrophage (Mg/Mp) markers are enriched only in the microglia/macrophages, regardless of Rbpms expression, and in the presumed RGC cells isolated by Li et al. (2022), but are not enriched in Tran et al. (2019) uninjured and injured RGCs or in axon-regenerating RGCs isolated by Jacobi et al. (2022).

The range of Rbpms expression in the Rbpms+ microglia/macrophages was equivalent to levels observed in uninjured and injured RGCs (Fig. 1A). Rbpms was expressed in the RGC-labeled cells from studies by Li et al.^5^, and Jacobi et al.^6^ (Fig. 1A). However, the other RGC-specific markers Slc17a6 and Sncg, and neuronal-specific marker Syn1, were not enriched in the RGC-labeled cells in Li et al.^5^, but were enriched in cells from study by Jacobi et al.^6^ (Fig. 1A-E). Neuronal-specific marker Tubb3 was not enriched in cells from either study (Fig. 1D). Only 17 cells, which represent 5.7% of all RGC-labeled cells that regenerated axons in the study by Li et al.^5^, expressed the appropriate RGC markers.

We then analyzed expression of microglia/macrophage markers, Iba1, Cd45, Cd14, Cd16, and Cx3cr1. We found that these markers were enriched in microglia/macrophages (regardless of whether they expressed Rbpms) and in the presumed RGC cells from the study by Li et al.^5^, but not in any of the RGC conditions, even after Pten KO in study by Jacobi et al.^6^ (Fig. 1F-J). After ONC, injury-activated microglia and infiltrating macrophages migrate within the optic nerve and into the retina. They phagocytize the tracer dye injected into the injured optic nerve, and some of them migrate into the retina within a day (thus, they are FACS’ed as dye+). Therefore, it is insufficient to rely on expression of only one pan-RGC marker to identify RGCs within a mixed population of scRNA-seq-profiled retinal cell types, as this can lead to inclusion of microglia/macrophages. Co-expression of pan-RGC markers (including Slc17a6 and Sncg) is necessary, and the inclusion of neuronal-specific markers (e.g., Tubb3 and Syn1) is preferred, in order to identify RGCs by scRNA-seq.

We also found that the neuroprotective genes, Anxa2 and Mpp1, found by Li et al.^5^, are highly expressed in the Rbpms+ microglia/macrophages (Fig. 2A-B), and along with Rbpms are also present in macroglia previously analyzed by bulkRNA-seq (see mouse microglia dataset at https://www.brainrnaseq.org)^7^. We also further confirmed that the presumed RGC cells from the study by Li et al.^5^ are microglia/macrophages, by analyzing global transcriptomes (and not only the canonical cell type markers) and finding that they correlated and clustered more strongly with the microglia/macrophage datasets than with other RGC datasets (Fig. 2C). Analysis of global transcriptomes also confirmed that Rbpms+ microglia/macrophages are near identical (by correlation and associated clustering) to the Rbpms-microglia/macrophages, and thus are not any other retinal cell type but are microglia/macrophages (Fig. 2C). Thus, inadvertently, Li et al.^5^ screened microglia/macrophage genes (instead of the genes enriched in the RGCs that regenerated axons after Pten KO), and found several neuroprotective hits, such as Anxa2 and Mpp1, which are expressed in both Rbpms+ microglia/macrophage and RGCs (Fig. 2A-B).

**Figure 2.**
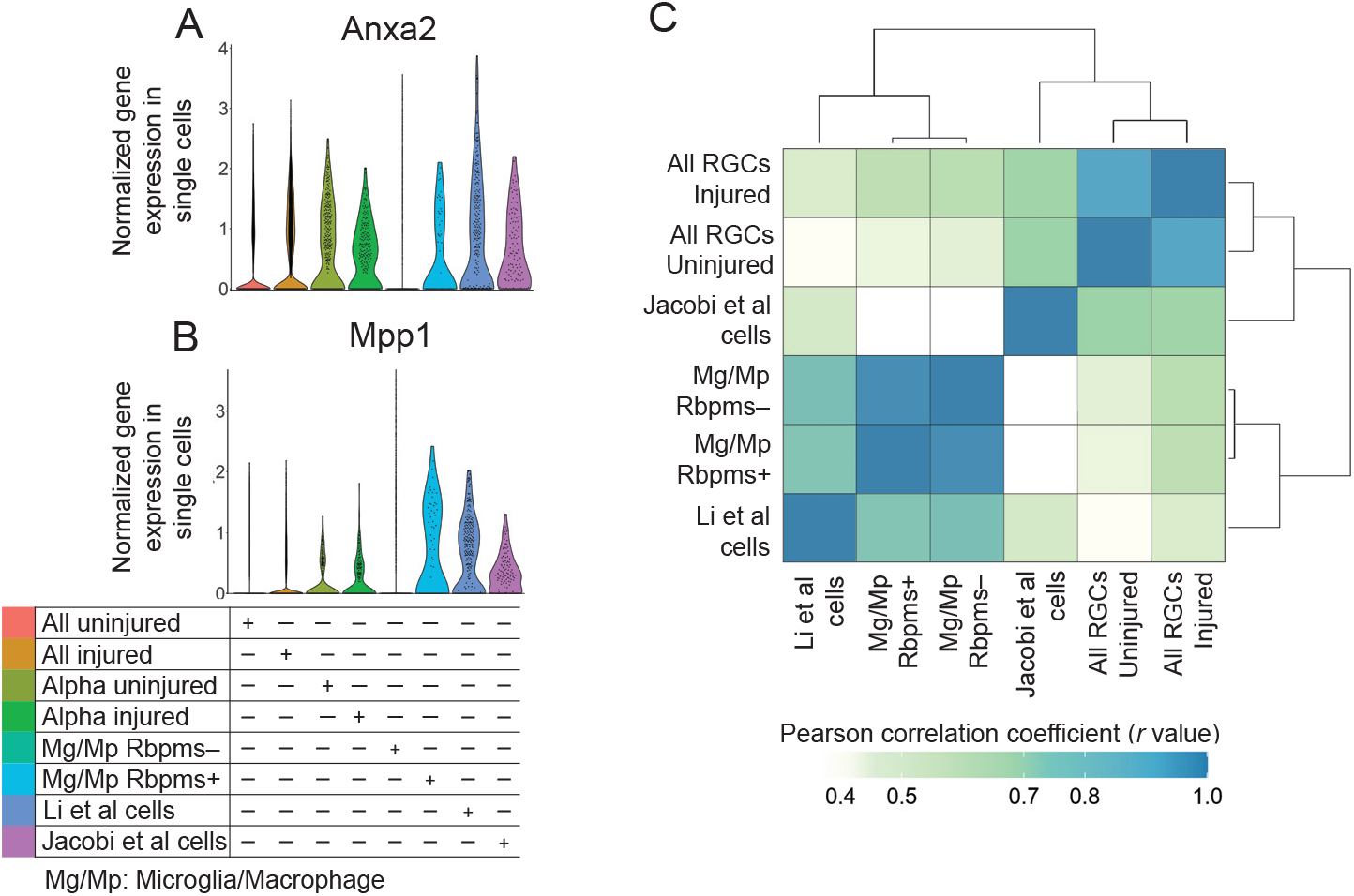
Violin plots of neuroprotective genes and global transcriptome analysis of single cells from different conditions. (**A**-**B**) Neuroprotective genes found in the screen by Li et al., Anxa2 (*A*) and Mpp1 (*B*), are highly expressed in the Rbpms+ microglia/macrophages, as well as in the Li et al. cells and in all RGC datasets. (**C**) Analysis of the global transcriptomes clustered and correlated presumed RGC cells from the study by Li et al. more strongly with the microglia/macrophage datasets than with other RGC datasets, whereas Jacobi et al. cells clustered and correlated more strongly with the RGC datasets than with microglia/macrophage datasets.

Because Li et al.^5^ found expression of certain αRGC markers in their cells, we analyzed whether they are expressed in microglia/macrophages too. The αRGC marker genes, Nefh (which encodes Neurofilament-H containing a non-phosphorylated epitope recognized by an SMI-32 antibody)^2,8^, Kcng4^2,9^, and Etv1^8,10^, were not enriched in cells from study by Li et al.^5^, with only a handful of cells expressing it within the 5.7% of cells that were indeed RGCs (Fig. 3A-C). However, Nefh and Kcng4 were enriched in the Jacobi et al.^6^ RGCs. Furthermore, although Kcng4 and Etv1 were downregulated in αRGCs after ONC, Kcng4 downregulation was prevented by Pten KO pre-treatment in Jacobi et al.^6^ cells (Fig. 3B-C). Other αRGC markers, Spp1^2,8,11^ and subtype C42-specific Fes^11^, however, were enriched in the Li et al.^5^ cells, but they were also enriched in the macrophages/microglia (Fig. 3D-E). Spp1 was also enriched in Jacobi et al.^6^ RGCs. Notably, Spp1 expression being substantially higher in the Li et al.^5^ cells compared to microglia/macrophages (Fig. 3E) is consistent with that Pten KO leads to Spp1 upregulation in some cell types^12-14^. Spp1 expression in macrophages was reported in prior studies as well^15-17^. Another αRGC subtype-specific marker, Kit (for C41)^11^, was not expressed except for in a few cells isolated by Li et al.^5^, and the remaining αRGC subtype-specific markers, Il1rapl2 (for C43) and Tpbg (for C45)^11^, were not enriched in the Li et al.^5^ cells. Thus, when using these markers for αRGCs in a mixed population of scRNA-seq-profiled cells, it is important to first evaluate for co-expression of pan-RGC markers.

**Figure 3.**
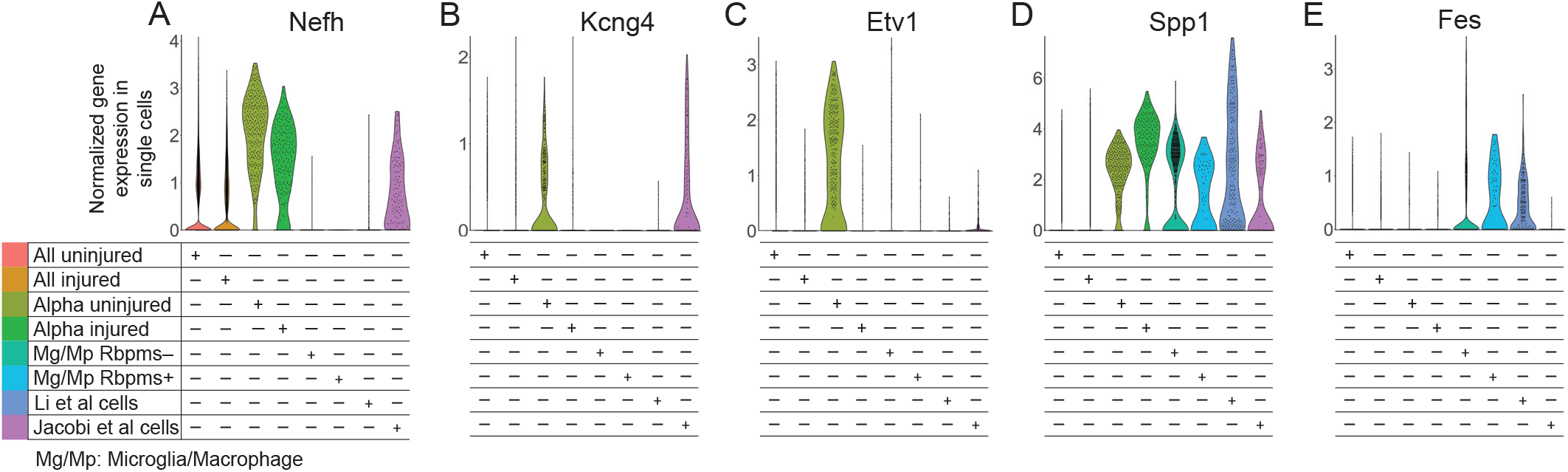
Violin plots of αRGC gene markers in single cells from different conditions. (**A**-**E**) Pan-αRGC markers Nefh (*A*), Kcng4 (*B*), and Spp1 (*D*), are enriched in Tran et al. (2019) αRGC subset and in axon-regenerating RGCs isolated by Jacobi et al. (2022). Spp1 (*D*) is enriched in the presumed RGC cells isolated by Li et al. (2022) more than in any other condition, but is also substantially enriched in the microglia/macrophages, regardless of Rbpms expression. Kcng4 (*B*) and Etv1 (*C*) αRGC markers are downregulated after ONC injury in Tran et al. (2019) αRGC subsets, but Kcng4 downregulation is prevented by Pten KO in axon-regenerating RGCs isolated by Jacobi et al. (2022). Fes (*E*) was enriched only in the presumed RGC cells isolated by Li et al. (2022) and in the microglia/macrophages.

## DISCUSSION

We reveled herein the existence of Rbpms+ and Spp1+ subpopulations of retinal microglia/macrophages, some of which also migrate into the retina from the optic nerve after activation by ONC injury. Rbpms is an established pan-RGC marker and Spp1 is an established αRGC type marker. Therefore, Rbpms+ and Spp1+ microglia/macrophages within retinal suspension could confound the identification of RGCs by scRNA-seq analysis, as occurred in a study by Li et al.^5^. However, serendipitous discovery of novel genes with neuroprotective potential by Li et al.^5^ study has significant implications for developing neuroprotective therapies, regardless of that microglia/macrophages analysis assisted in their identification.

We also show that in order to circumvent the potential pitfall of mistaking Rbpms+ microglia/macrophages for RGCs in scRNA-seq studies, detection of co-expression of pan-RGC markers (including Rbpms, Slc17a6, and Sncg) is necessary, and the inclusion of neuronal-specific markers (e.g., Tubb3 and Syn1) is preferred. Similarly, in order to circumvent the potential pitfall of mistaking Spp1+ microglia/macrophages for αRGCs in scRNA-seq studies, it is important to first evaluate for co-expression of pan-RGC markers other than just Rbpms, and the inclusion of other pan-αRGC markers (e.g., Nefh and Kcng4) is preferred, with the caveat that Kcng4 is downregulated after ONC injury unless pre-treated (with Pten KO in this case). Similar considerations need to be made when relying on αRGC subtype C42 marker Fes, as it is also expressed in a subpopulation of Rbpms+ microglia/macrophages.

## MATERIALS AND METHODS

The cells identified as RGCs, which regenerated axons in response to Pten knockout, were processed for scRNA-seq by plate-based SmartSeq2 in studies by Li et al.^5^ and Jacobi et al.^6^. ScRNA-seq counts-matrix were assembled following alignment to the mouse genome and transcriptome using the Hisat2-Cufflinks pipeline. Counts-matrix for Li et al.^5^ (GSE206626) and Jacobi et al.^6^ (GSE202155) were obtained from Gene Expression Omnibus. RGCs were selected per criteria specified in the respective paper’s methods sections. In scRNA-seq dataset from Li et al.^5^, 297 cells were initially identified as presumed RGCs (that regenerated axons), but only 17 cells were left after exclusion of macrophages/microglia in retrospect analysis reported herein (the other group of RGC-labeled cells in Li et al.^5^ study that survived but did not regenerate axons had just 8 RGCs and the remaining cells were microglia/macrophages when the appropriate markers were used). In the scRNA-seq dataset from Jacobi et al.^6^, 120 cells were identified as RGCs. 10x Genomics 3’-droplet based scRNA-seq dataset of adult uninjured and injured (2 weeks after ONC) RGCs, which included αRGCs, was obtained from Tran et al.^11^ (GSE137400). We generated 10x Genomics 3’-droplet based scRNA-seq dataset of macrophage/microglia from retina/optic nerve, and demultiplexing and alignment to the mouse genome and transcriptome was performed using the CellRanger pipeline. Comparative analysis of scRNA-seq datasets by Violin plots was performed using the R package Seurat v4.3.0. Correlation matrix was generated using R software^18^. The macrophage/microglia scRNA-seq data, and the integrated data from different cell types and conditions, which we generated, are available through the NCBI GEO accession *Pending*.

## AUTHOR CONTRIBUTIONS

W.C.T performed the analysis. E.F.T. wrote the manuscript.

## ACKNOWLEDGMENTS

This work was supported by grants from The University of Connecticut School of Medicine, Start-Up Funds (to E.F.T.), and the National Institutes of Health (NIH) (Grant R01-EY029739, to E.F.T.). Portions of this research were conducted at the High Performance Computing Facility, University of Connecticut.

## REFERENCES

1. Park KK, Liu K, Hu Y, et al. Promoting axon regeneration in the adult CNS by modulation of the PTEN/mTOR pathway. Science. Nov 7 2008;322(5903):963–6.

2. Duan X, Qiao M, Bei F, Kim IJ, He Z, Sanes JR. Subtype-Specific Regeneration of Retinal Ganglion Cells following Axotomy: Effects of Osteopontin and mTOR Signaling. Neuron. Mar 2015;85(6):1244–56. doi:10.1016/j.neuron.2015.02.017

3. Kim J, Sajid MS, Trakhtenberg EF. The extent of extra-axonal tissue damage determines the levels of CSPG upregulation and the success of experimental axon regeneration in the CNS. Sci Rep. Jun 2018;8(1):9839. doi:10.1038/s41598-018-28209-z

4. Yungher BJ, Luo X, Salgueiro Y, Blackmore MG, Park KK. Viral vector-based improvement of optic nerve regeneration: characterization of individual axons’ growth patterns and synaptogenesis in a visual target. Gene Ther. Oct 2015;22(10):811–21. doi:10.1038/gt.2015.51

5. Li L, Fang F, Feng X, et al. Single-cell transcriptome analysis of regenerating RGCs reveals potent glaucoma neural repair genes. Neuron. 08 17 2022;110(16):2646-2663.e6. doi:10.1016/j.neuron.2022.06.022

6. Jacobi A, Tran NM, Yan W, et al. Overlapping transcriptional programs promote survival and axonal regeneration of injured retinal ganglion cells. Neuron. 08 17 2022;110(16):2625-2645.e7. doi:10.1016/j.neuron.2022.06.002

7. Bennett ML, Bennett FC, Liddelow SA, et al. New tools for studying microglia in the mouse and human CNS. Proc Natl Acad Sci U S A. Mar 22 2016;113(12):E1738–46. doi:10.1073/pnas.1525528113

8. Rheaume BA, Jereen A, Bolisetty M, et al. Single cell transcriptome profiling of retinal ganglion cells identifies cellular subtypes. Nat Commun. Jul 2018;9(1):2759. doi:10.1038/s41467-018-05134-3

9. Krieger B, Qiao M, Rousso DL, Sanes JR, Meister M. Four alpha ganglion cell types in mouse retina: Function, structure, and molecular signatures. PLoS One. 2017;12(7):e0180091. doi:10.1371/journal.pone.0180091

10. Martersteck EM, Hirokawa KE, Evarts M, et al. Diverse Central Projection Patterns of Retinal Ganglion Cells. Cell Rep. Feb 2017;18(8):2058–2072. doi:10.1016/j.celrep.2017.01.075

11. Tran NM, Shekhar K, Whitney IE, et al. Single-Cell Profiles of Retinal Ganglion Cells Differing in Resilience to Injury Reveal Neuroprotective Genes. Neuron. 12 2019;104(6):1039-1055.e12. doi:10.1016/j.neuron.2019.11.006

12. Kim HJ, Ryu J, Woo HM, et al. Patterns of gene expression associated with Pten deficiency in the developing inner ear. PLoS One. 2014;9(6):e97544. doi:10.1371/journal.pone.0097544

13. Sato W, Horie Y, Kataoka E, et al. Hepatic gene expression in hepatocyte-specific Pten deficient mice showing steatohepatitis without ethanol challenge. Hepatol Res. Apr 2006;34(4):256–65. doi:10.1016/j.hepres.2006.01.003

14. Lu TL, Huang YF, You LR, et al. Conditionally ablated Pten in prostate basal cells promotes basal-to-luminal differentiation and causes invasive prostate cancer in mice. Am J Pathol. Mar 2013;182(3):975–91. doi:10.1016/j.ajpath.2012.11.025

15. Zhang Y, Du W, Chen Z, Xiang C. Upregulation of PD-L1 by SPP1 mediates macrophage polarization and facilitates immune escape in lung adenocarcinoma. Exp Cell Res. Oct 15 2017;359(2):449–457. doi:10.1016/j.yexcr.2017.08.028

16. Dong B, Wu C, Huang L, Qi Y. Macrophage-Related SPP1 as a Potential Biomarker for Early Lymph Node Metastasis in Lung Adenocarcinoma. Front Cell Dev Biol. 2021;9:739358. doi:10.3389/fcell.2021.739358

17. Morse C, Tabib T, Sembrat J, et al. Proliferating SPP1/MERTK-expressing macrophages in idiopathic pulmonary fibrosis. Eur Respir J. Aug 2019;54(2)doi:10.1183/13993003.02441-2018

18. R-Core_Team. R: A language and environment for statistical computing. R Foundation for Statistical Computing. http://www.R-project.org; 2014.

